# Connectomes for 40,000 UK Biobank participants: A multi-modal, multi-scale brain network resource

**DOI:** 10.1101/2023.03.10.532036

**Authors:** L. Sina Mansour, Maria A. Di Biase, Robert E. Smith, Andrew Zalesky, Caio Seguin

## Abstract

We mapped functional and structural brain networks for more than 40,000 UK Biobank participants. Structural connectivity was estimated with tractography and diffusion MRI. Resting-state functional MRI was used to infer regional functional connectivity. We provide high-quality structural and functional connectomes for multiple parcellation granularities, several alternative measures of interregional connectivity, and a variety of common data pre-processing techniques, yielding more than one million connectomes in total and requiring more than 200,000 hours of compute time. For a single subject, we provide 28 out-of-the-box versions of structural and functional brain networks, allowing users to select, e.g., the parcellation and connectivity measure that best suit their research goals. Furthermore, we provide code and intermediate data for the time-efficient reconstruction of more than 1,000 different versions of a subject’s connectome based on an array of methodological choices. All connectomes are available via the UK Biobank data sharing platform and our connectome mapping pipelines are openly available. In this report, we describe our connectome resource in detail for users, outline key considerations in developing an efficient pipeline to map an unprecedented number of connectomes, and report on the quality control procedures that were completed to ensure connectome reliability and accuracy. We demonstrate that our structural and functional connectivity matrices meet a number of quality control checks and replicate previously established findings in network neuroscience. We envisage that our resource will enable new studies of the human connectome in health, disease and aging at an unprecedented scale.

## Background & Summary

Different aspects of brain connectivity can be quantified using different MRI modalities: diffusion-weighted MRI data can be utilized to map structural brain networks of white-matter connections^1,2^; alternatively, functional MRI data can be used to map functional connectivity networks describing inter-regional interactions in brain activity^3,4^. These network representations of brain connectivity are referred to as connectomes^5,6^. Establishing a large-scale community biobank of structural and functional human connectomes will enable a diverse range of research into brain networks in health and disease.

The importance of large-scale neuroimaging biobanks is increasingly recognized as key to addressing reproducibility concerns in neuroscience^7–12^. The UK biobank (UKB)—a population study containing in-depth biomedical, health, and environmental data—is the world’s largest neuroimaging resource (with ∼ 45,000 imaging sessions acquired from ~40,000 participants thus far)^8,13–15^. This biobank offers tremendous potential for research on early diesase prediction and alignment of image-derived phenotypes (IDPs) with cognitive, behavioral, genetic, and medical observations. The availability of longitudinal neuroimaging data accompanying constantly updated clinical records enables prospective neuroscientific research at a population scale. To facilitate this effort, the UKB has released a range of important quantitative neuroimaging derivatives, including regional measures of brain structure, microstructure, and function. At present however, measures of brain *connectivity* are not a part of this resource. Mapping connectomes from neuroimaging data at scale is computationally burdensome and requires significant technical expertise. Establishing a brain connectivity biobank for the UKB will ensure rapid access to connectomes for researchers without expertise or computing resources for large-scale connectivity mapping, facilitate reproducible neuroscience practices, enhance UKB utilization among the research community and ultimately lead to new discoveries about brain networks.

Here, we introduce our novel human connectome resource of brain atlases and connectivity matrices mapped for more than 40,000 adults participating in the UKB (see Figure 1). We provide functional activity time series and structural connectivity matrices for multiple parcellation schemes, several alternative measures of interregional connectivity and a variety of common data pre-processing techniques, yielding 27 brain-wide functional time series (enabling flexible and efficient access to various functional connectivity metrics), and 28 structural connectomes per imaging session, and more than one million connectomes and time-series in total. This required development of highly efficient connectome mapping pipelines and storage formats. Connectivity data is made available in compact and easy-to-use data formats and our connectome mapping pipelines are openly available. We completed extensive quality control procedures to ensure the accuracy and reliability of all connectivity matrices. This report aims to describe our connectome resource in detail for prospective users, provide insight into the key considerations that shaped the development of our connectome mapping pipeline and outline quality control procedures. We envisage that this resource will be of high utility and complement the current IDPs available in the UKB.

**Figure 1.**
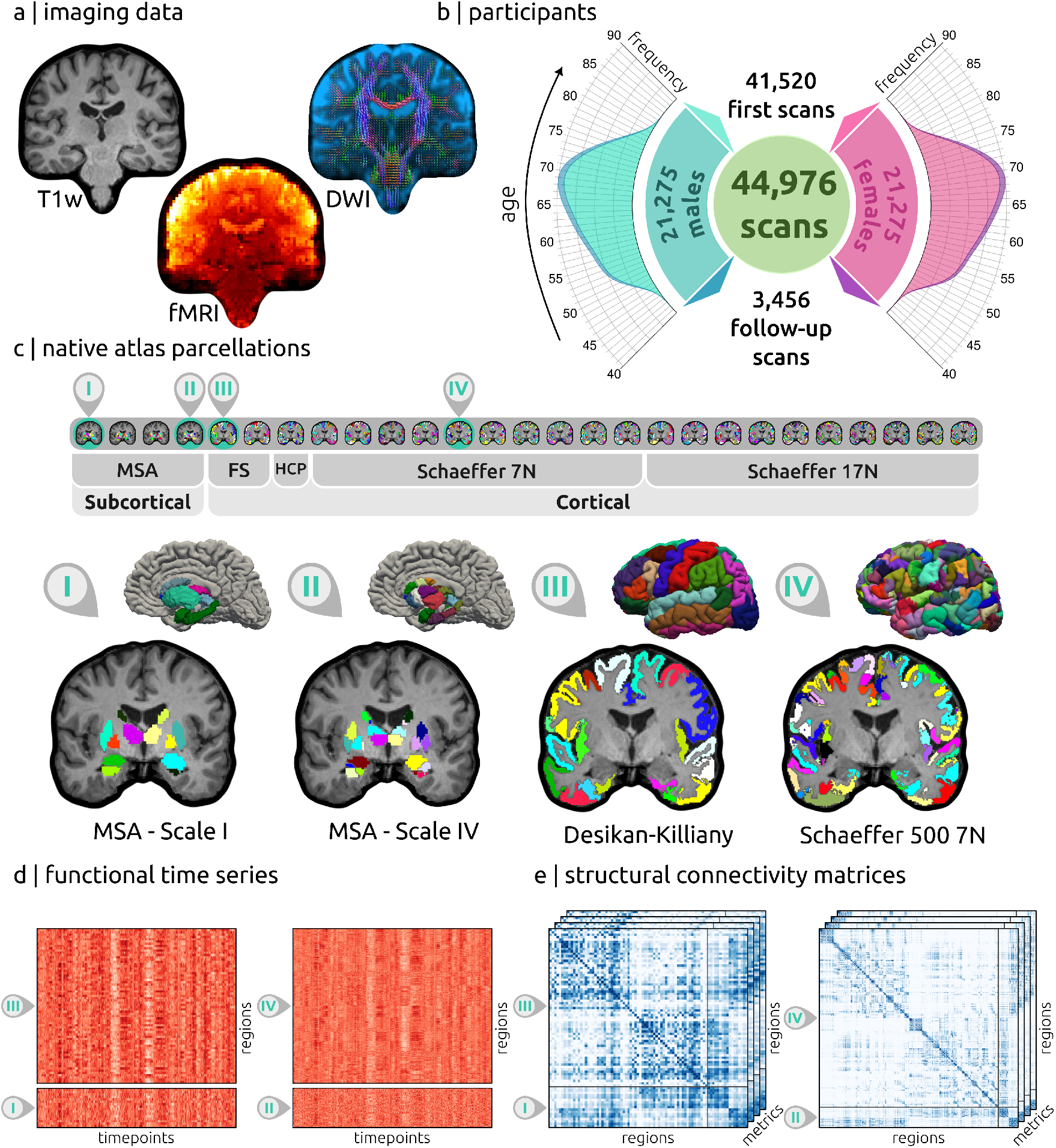
A comprehensive connectome biobank for the UK Biobank. (a) Structural and functional connectomes were mapped from diffusion and resting-state functional MRI data, respectively, while T1-weighted structural MRI was used for brain alignment. (b) Distribution of age and sex in the 41,520 UK Biobank participants with neuroimaging data available. Longitudinal data was available for 3,456 participants. (c) Various whole-brain atlases were used to map individual parcellations in native volumetric space. A total of 27 alternative parcellation schemes were computed. Four representative subcortical (I and II) and cortical (III and IV) parcellations of different granularities are illustrated. (d) Resting-state functional time series and (e) various structural connectivity matrices were computed for different parcellations. The time series and structural connectivity matrices are depicted for the same four sample parcellations. Abbreviations; MSA: Melbourne Subcortical Atlas, FS: FreeSurfer atlases, HCP: HCP-MMP1.0 atlas.

## Methods

### UK Biobank data

#### Background

The UKB is a large-scale dataset comprising over 500,000 participants (aged 40-69 years). The biobank is publicly available to advance health-related research^13^. Importantly, access to full health records (which are updated over time), and a wide range of longitudinal, long-term clinical, phenotypic, and genomic data is available to researchers^15^. In addition, UKB aims to acquire longitudinal neuroimaging data for 100,000 participants, with ∼ 45,000 separate MRI sessions for ∼ 40,000 participants released thus far^8,14^.

#### Participants

Brain MRI data sourced for the connectome biobank consists of 44,976 separate imaging sessions (23,701 females and 21,275 males) from 41,520 individuals (21,951 females and 19,569 males) aged 44–82 years (*µ* = 64.0, *σ* = 7.6 years) at time of acquisition. For 3,456 individuals (1,750 females and 1,706 males) a single longitudinal follow-up MRI session was acquired 1–7 years (*µ* = 2.35, *σ* = 0.71 years) after the first acquisition. Further details regarding recruitment protocols are provided elsewhere^14^. Imaging sessions that did not pass the existing UKB preprocessing and quality control pipeline^8^ were excluded.

#### MRI data acquisition and existing preprocessing

A detailed description of the MRI data acquisition and preprocessing pipeline is provided elsewhere^8^. All modalities were acquired on 3T Siemens Skyra scanners using the standard Siemens 32-channel head coil. The T1-weighted structural brain images (Data-Field 20252) were acquired using a 3D MPRAGE acquisition at 1mm isotropic resolution with a 256mm superior-inferior field of view^8^. The preprocessing steps included gradient distortion correction (GDC)^16,17^, skull stripping^18^, linear and nonlinear registration to MNI152 standard space^19–21^, and defacing^8^. Additional derivative data precalculated and provided by the UKB resource include macroscopic tissue segmentation with FSL FAST^22^, subcortical modeling with FSL FIRST^23^, and brain segmentation including cortical surface estimation with Freesurfer^24^.

Resting-state BOLD data (Data-Field 25751) were acquired with a multi-band gradient echo EPI sequence^25–27^, with an acquisition time of ∼ 6 minutes, for a total of 490 volumes, with a spatial resolution of 2.4mm isotropic voxels (TE/TR=39/735 ms, MB=8, no in-plane acceleration, flip angle 52^*°*^, conventional fat saturation)^14^. Preprocessing steps^8^ consisted of the FSL MELODIC pipeline^28^ (EPI susceptibility distortion correction, GDC, motion correction with FSL MCFLIRT^20^, grand-mean intensity normalization, and high-pass temporal filtering) followed by an ICA + FIX step to suppress remaining artifact components^29,30^.

Diffusion-weighted MRI (dMRI) data (Data-Field 20250) were acquired using a multi-band spin echo EPI sequence^31^, with an acquisition time of ∼ 7 minutes, with 100 unique diffusion sensitisation directions distributed equally across two shells (*b*-values: 1000, 2000 *s/mm*^2^), and 5 *b*=0 volumes, with a spatial resolution of 2mm isotropic voxels (MB=3, no in-plane acceleration, TE/TR=92/3600 ms, partial Fourier 6/8, conventional fat saturation). 3 additional *b*=0 volumes were acquired with reversed phase encoding direction to enable susceptibility field estimation^32^. The data were preprocessed with a pipeline^8^ consisting of correction for eddy current and head motion^33–35^ followed by GDC^36^. Additionally, FSL’s `dtifit`^37^ and the NODDI toolbox^38^ were used to generate voxelwise microstructural parameters: fractional anisotropy (FA), tensor mode (MO), mean diffusivity (MD), *b*=0 signal intensity (S0), intra-cellular volume fraction (ICVF), isotropic volume fraction (ISOVF), and orientation dispersion index (ODI).

### Connectomic nomenclature

Table 1 provides brief definitions for terms commonly used in this paper.

**Table 1.**
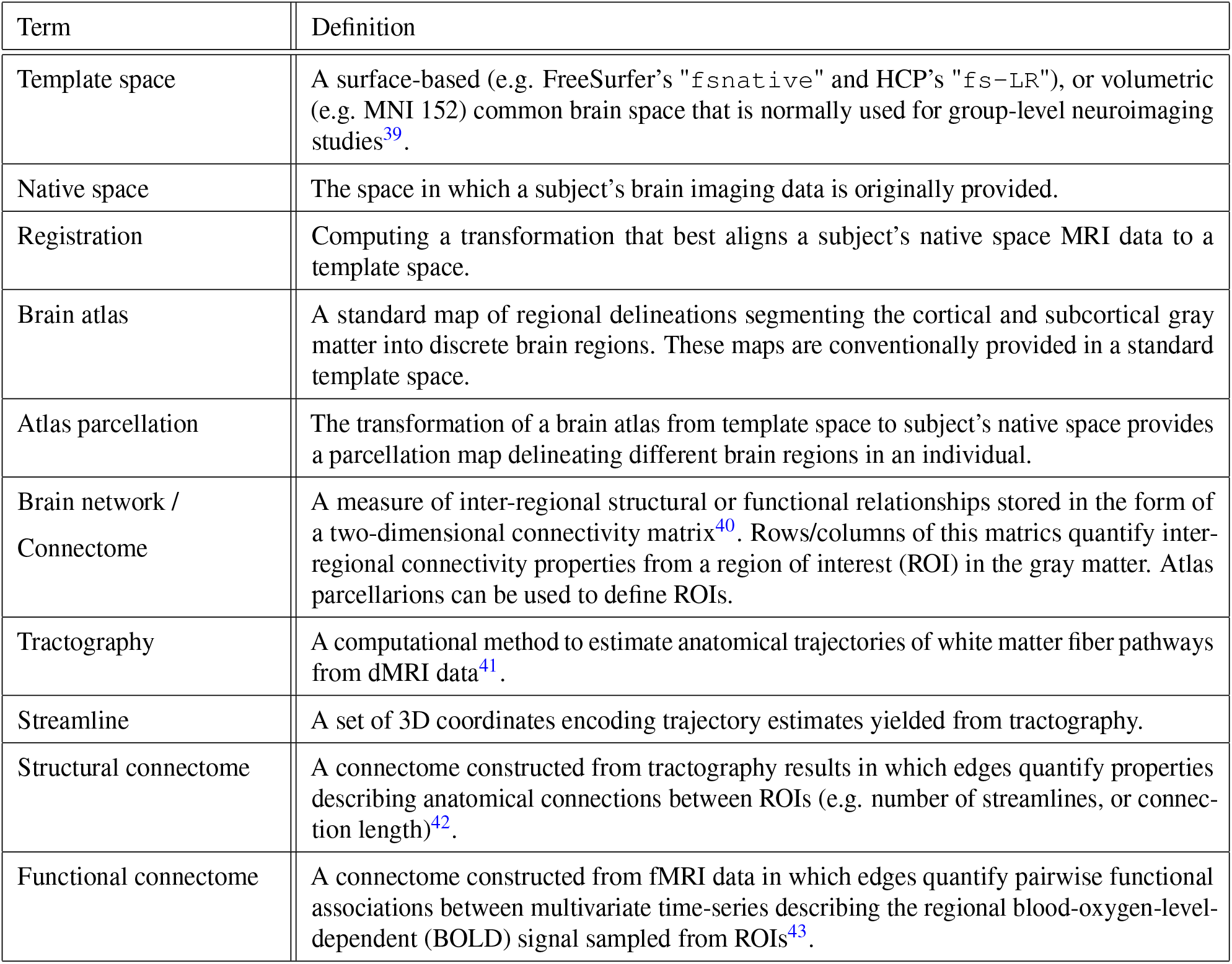
A description of frequently used terms.

### Connectome reconstruction pipelines

As detailed below, we developed novel connectome reconstruction pipelines to satisfy five competing demands:

1. *Connectome quality*, producing high-quality brain networks by using state-of-the-art pipelines to infer interregional brain connectivity from diffusion and functional neuroimaging data.
2. *Accessibility*, facilitating streamlined access to connectome data in an efficient and easy to use format.
3. *Flexibility*, enabling users to choose from connectomes mapped using a broad range of methodological preferences.
4. *Computational requirements*, as our pipeline needed to be executed on data from ∼ 45,000 MRI sessions, which is a considerable burden even when making use of high-performance computing (HPC) services.
5. *Storage requirements*, as the generated connectivty data needs to be stored both in the short term on local storage during calculation, and in the long term on UKB storage infrastructure.

Hence, the pipeline was developed to provide a good balance between these competing aims and prepare a comprehensive, versatile, and high-quality brain connectivity resource. This included the development of new utilities in software packages used to map conenctomes (e.g., *MRtrix3*^44^) with an explicit focus on reducing computational and storage requirements.

A crucial conflict identified within this set of demands lies at the intersection of accessibility, flexibility, and storage requirements. On the one hand, user flexibility is boosted by providing many versions of the same connectome, mapped according to different methodological choices, such as different atlases or interregional connectivity measures. On the other hand, catering for flexibility leads to a combinatorial increase in the number of connectomes, which would be prohibitively expensive to store and access for all UKB subjects. This would be particularly the case for SC matrices, for which there are many options for the metric of connectivity.

We addressed this important issue as follows. For a subset of connectome configurations, deemed to be of broad applicability, connectivity matrices were calculated and uploaded to the UKB for direct user access. Where an alternative configuration is desired, we provide both the requisite intermediate data (ie. for which the most expensive computational processes have already been performed) and a software tool that uses these data to efficiently calculate the connectome of interest, such that any combination of connectome attributes can be chosen. This tool, along with the relevant code for the pipeline in its entirety, is accessible from github.com/sina-mansour/UKB-connectomics.

An overview flowchart of the pipeline is presented in Figure 2. In short, a set of automated pipelines were implemented to perform three main tasks:

**Figure 2.**
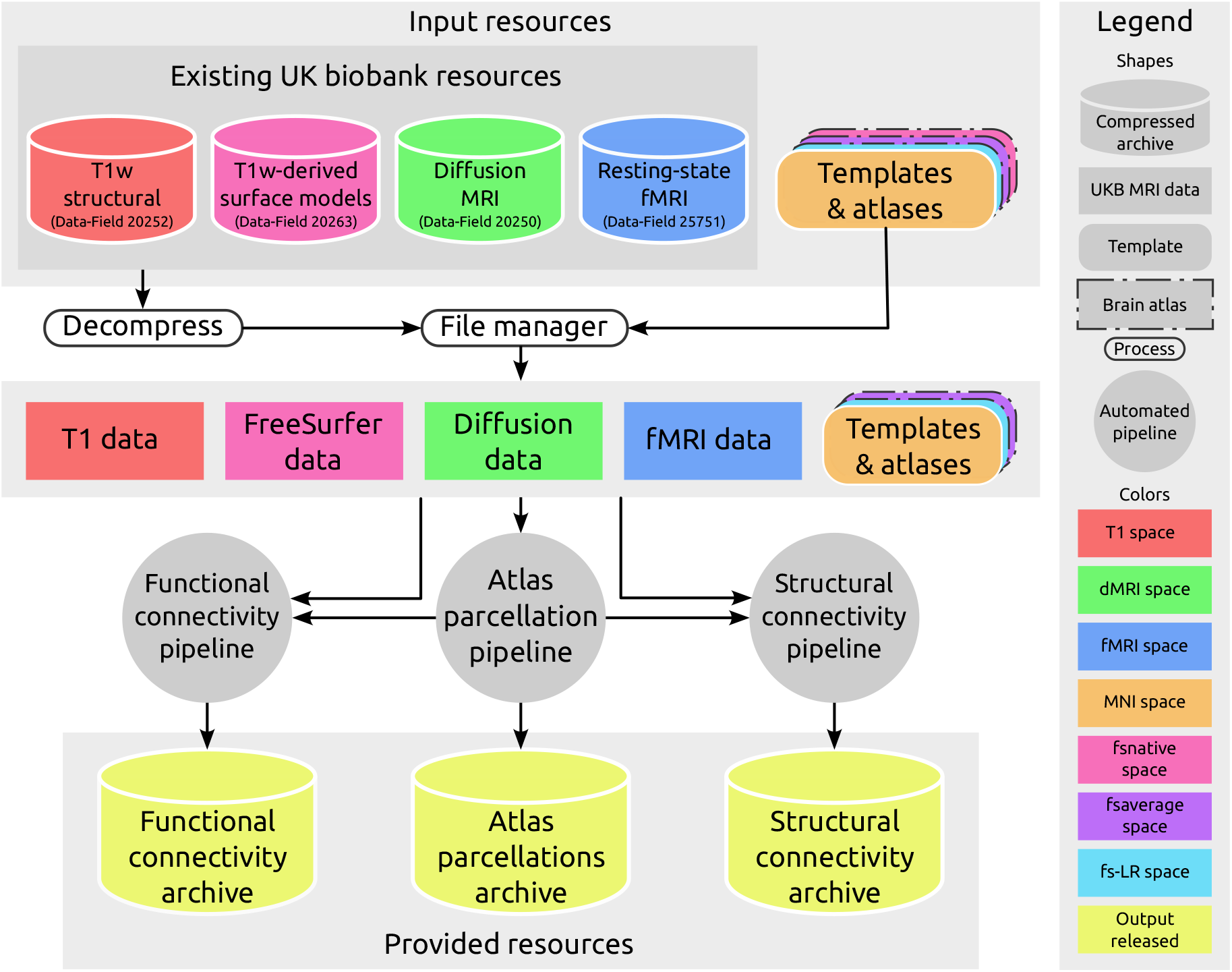
Schematic flowchart of the complete pipeline. Compressed bulk data archives from UKB along with publicly available brain parcellation atlases and templates were used as inputs of the connectivity mapping pipelines. These inputs were used in three separate steps to generate parcellations in subject space, that were subsequently utilized in functional and structural connectome construction pipelines. Flowcharts of these steps are detailed in the ensuing figures. For every imaging session, the outputs of these three steps are provided as separate compressed archives.

1. *Atlas parcellations*: Generate volumetric atlas parcellations in subject space, segmenting cortical and subcortical gray matter into distinct brain regions
2. *Functional connectivity*: Derive resting-state functional time series of brain activity within these parcels to enable functional connectivity (FC) estimation
3. *Structural connectivity*: Estimate white matter fiber orientations from diffusion MRI data and perform tractography to map structural connectivity (SC) matrices

The following sections provide a detailed explanation of every step in the connectome reconstruction pipeline.

### Brain atlas parcellations

For parcellation of the cortical and subcortical gray matter into distinct nodes of a connected network, here we focus strictly on the conventional approach in the domain of neuroimaging connectomics, where spatial correspondence is established between a pre-generated atlas defined in some template space and the subject-specific T1-weighted image (see Table1)^40,42^. Importantly, it is well established that the choice of atlas, and in particular the granularity of gray matter segmentations, may impact findings in brain connectivity studies^45,46^. To address this issue, as shown in Figure 3, we consider a total of 23 cortical and 4 subcortical atlases. This provides researchers with the ability to choose the most appropriate parcellation for their study, investigate connectivity features at multiple spatial scales, and replicate analyses across different parcellation schemes.

**Figure 3.**
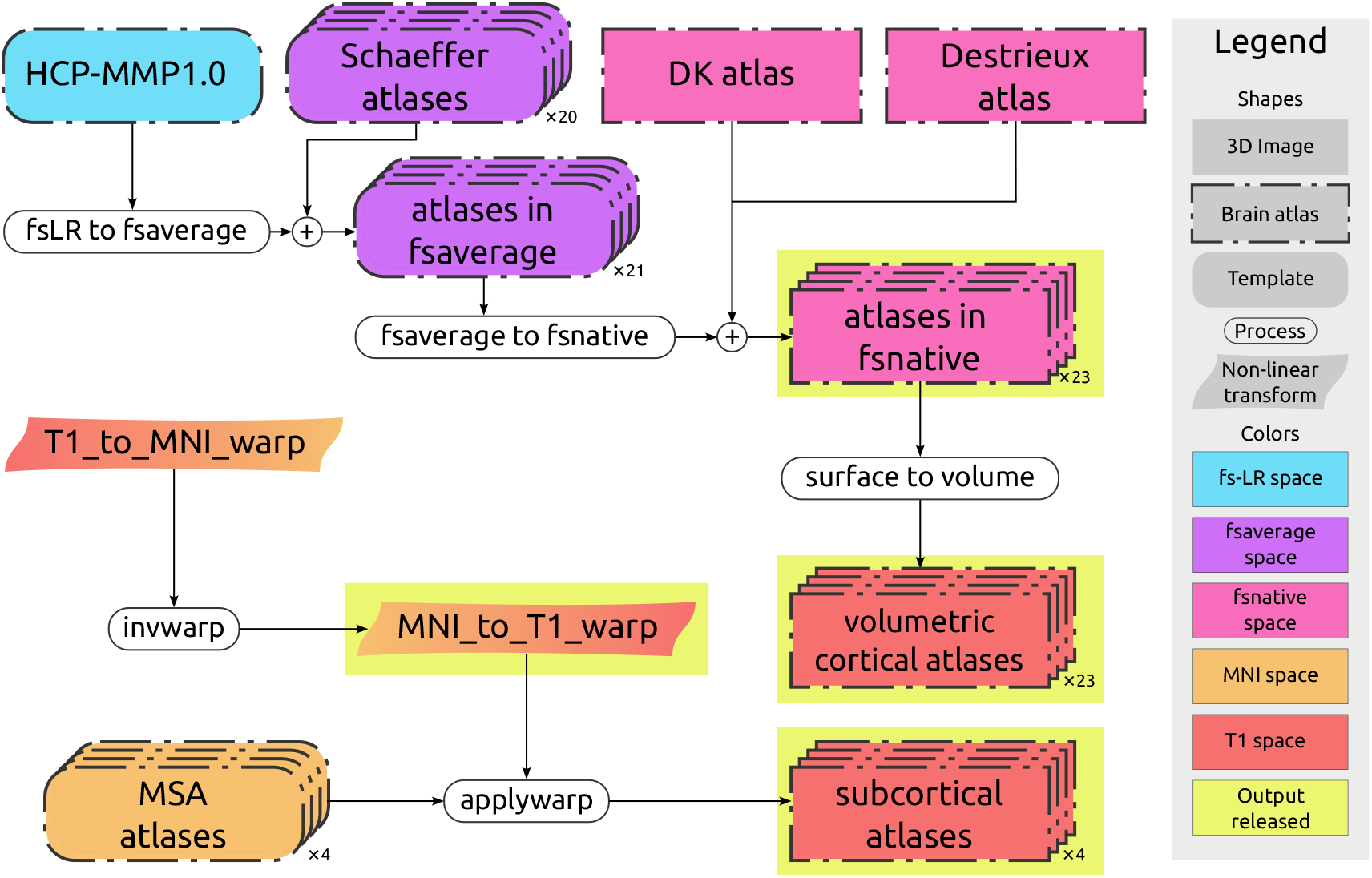
Flowchart for the brain atlas parcellation pipeline. A total of 23 cortical and 4 subcortical atlases, transformed from their respective template spaces, were mapped to each individual’s native volumetric space. These volumetric atlas parcellations, as well as additional supporting files (volumetric warp and native surface delineations of cortical atlases), are made available in the atlas parcellations compressed archive (indicated with yellow highlight).

An important detail of our handling of parcellation data is that cortical and subcortical gray matter parcellations were processed entirely independently of one another, and are only combined immediately prior to connectome construction. This permits independent selection of both cortical and subcortical parcellations for any given connectome configuration.

We computed volumetric brain atlases in every individual’s native space for a total of 23 cortical and 4 subcortical atlases of the human brain. The cortical atlases included 20 different Scheaffer parcellations^47^ (derived from two different sets of functional networks^48^ and sub-divided into different parcellation granularities ranging 100–1000 nodes), the Human Connectome Project’s multimodal parcellation^49^ (HCP-MMP1.0; also known as the Glasser atlas), the Desikan-Killiany atlas^50^ (FreeSurfer’s “`aparc`”), and the Destrieux Atlas^51^ (FreeSurfer’s “`aparc.a2009s`”). The subcortical atlases included 4 different spatial scales of the Melbourne Subcortical Atlas (MSA)^52^.

All atlases were transformed to each individual’s native volumetric space. For the subcortical atlases, the UKB provides non-linear warp files from the subject’s T1-weighted image to the MNI152 template^21^, which is the space in which the MSA parcellations are defined. We first computed the inverse of this warp (ie. from MNI152 to the subject’s T1-weighted image), then applied this transformation to each of the four parcellations using nearest-neighbor interpolation (FSL’s “`applywarp`”). To map subject-specific cortical atlases, we started from surface-based labels provided in different template spaces^47,49–51^. For each participant, we aligned these labels to the subject’s native cortical surface representation (FreeSurfer’s “`fsnative`”), as this provides superior anatomical accuracy compared to volumetric registrations. To transform surface labels into the final volumetric atlases, we first defined, for each subject, a cortical ribbon at the interface between gray and white matter. Lastly, we assigned every voxel in the ribbon the label of its nearest surface vertex. The surface transformations made use of in-house methods incorporating various functionalities from the Connectome Workbench^53^, FreeSurfer^24^, and NiBabel^54^. All parcellations in subject space (both surface and volumetric representations), the inverse nonlinear MNI warp, and associated conversion scripts (to expedite use of atlases not processed here) are made available (see *Data Records and Code Availability*). Additional scripts are also provided to combine cortical and subcortical atlases, which are required for mapping SC matrices comprising combinations of cortical and subcortical parcellations beyond the ones readily provided in our resource (see *Structural connectivity: matrices* section for detail).

In some circumstances, both cortical and subcortical sources may attribute a parcel to the same voxel in native space. This occurred most predominantly in the hippocampal region when using the HCP-MMP 1.0 / Glasser atlas. The software used for connectome construction and provided to the research community prioritizes cortical labels where this occurs.

### Functional connectivity

FC characterizes statistical dependencies between the BOLD time series recorded from different brain regions^40^. Several computational approaches can be used to map functional connectivity from BOLD signals^4,55^. Here, we provide regionally averaged BOLD time series for the same brain parcellations atlases used to map SC matrices. This enables researchers to easily compute FC matrices using their favorite methods, and also allows for analyses of dynamic and time-varying FC. The flowchart of FC processing is shown in Figure 4. All parcellations were first resampled from the subject’s T1-weighted image voxel grid to the subject’s preprocessed fMRI voxel grid (FreeSurfer’s `mri_vol2vol`). For each parcel, the mean time series across the fMRI voxels ascribed to that region was calculated. By providing these parcellated BOLD time series, we enable flexibility and convenience in mapping functional connectivity across a wide range of alternative spatial resolutions and methodological approaches. For instance, FC derived from statistical correlation-based measures (e.g. Pearson’s r) can be directly computed from the time series data in a few milliseconds.

**Figure 4.**
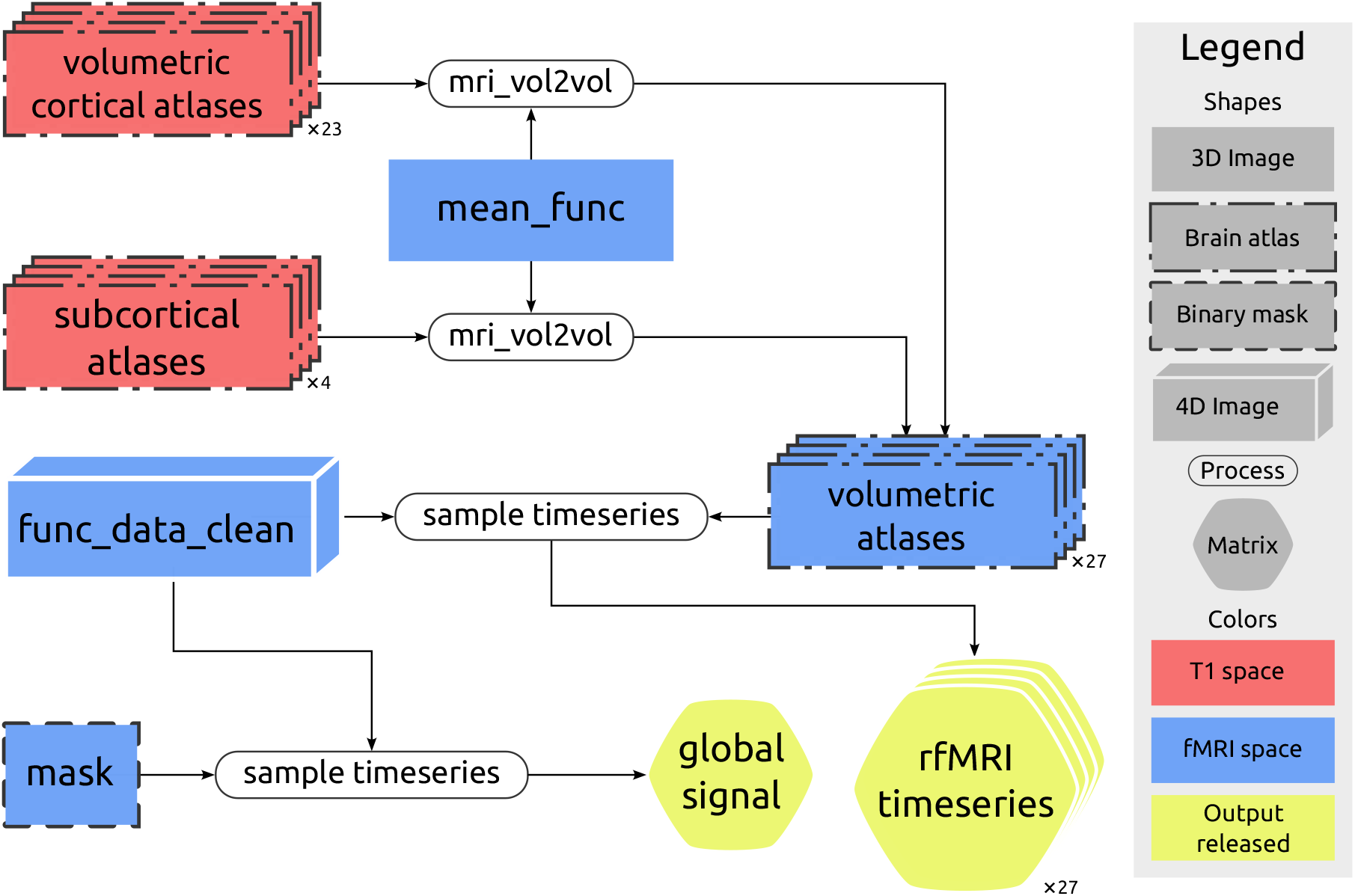
Flowchart for the functional connectivity mapping pipeline. BOLD regional time series are provided for 27 brain parcellation atlases, enabling rapid computation of functional connectivity matrices. The global signal time series was additionally computed. Note: yellow color indicates output resources that are made available in the functional connectivity compressed archive.

Global signal regression (GSR) is an fMRI preprocessing technique with potential merits and drawbacks that are subject to debate^56–64^. We thus calculated the global signal time series and provide those data separately to provide researchers with the flexibility to perform GSR if desired. The global signal was computed by averaging the BOLD time series over all voxels belonging to the anatomical brain mask.

### Structural connectivity: tractography

Structural connectomes provide a network representation of the brain white matter axonal architecture^40^. Whole-brain tractography is used to map white matter axonal pathways from an individual’s diffusion MRI data and enable estimation of the connectivity properties of these pathways using any of a number of structural connectivity measures. Figure 5 shows how whole-brain tractograms were computed using probabilistic tractography as implemented in *MRtrix3*^44^.

**Figure 5.**
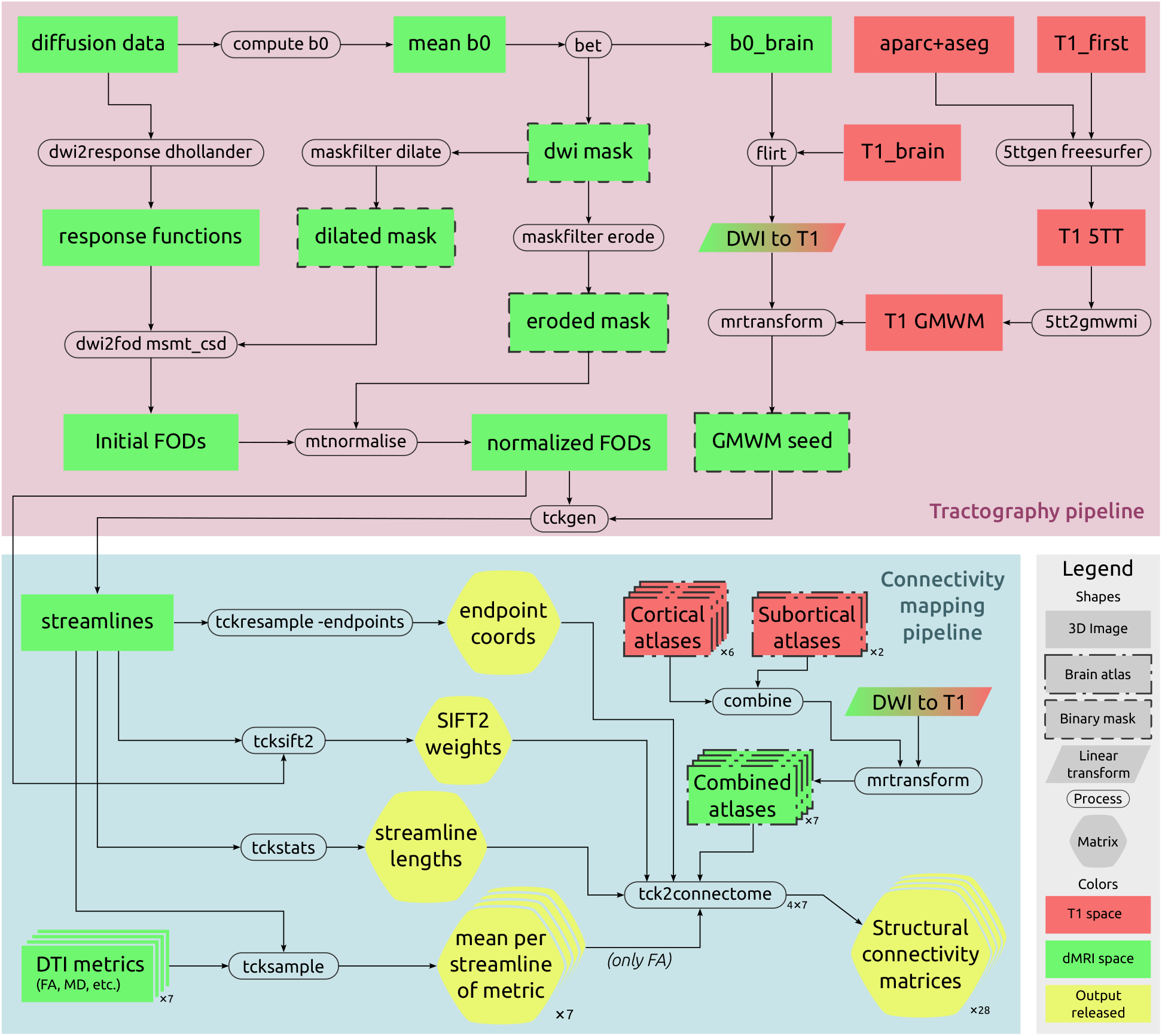
Flowchart for the structural connectivity mapping pipeline. The pipeline is divided into two sections. First, white matter tractography is used to generate streamlines from diffusion-weighted imaging data. Next, these streamlines are used to estimate interregional connectivity properties based on various measures of structural connectivity. The streamline endpoint coordinates, connectivity metrics per streamline, and the subset of 28 connectome matrix configurations that were explicitly generated, are all provided in the structural connectivity compressed archive (shown in yellow).

A brain mask was derived by skull-stripping the mean of DWI volumes without any diffusion weighting (*b*=0 volumes) and executing the FSL `bet` tool with parameters tuned for diffusion-weighted imaging (DWI) data^18^. In comparison to the DWI masks provided by the UKB, these masks were deemed more accurate and resulted in better registration between the T1 and DWI spaces. Macroscopic tissue response functions^65^ for white matter (WM), gray matter (GM), and cerebrospinal fluid (CSF) were estimated with an unsupervised heuristic^66,67^. Multi-shell, multi-tissue (MSMT) constrained spherical deconvolution (CSD)^68^ was used to estimate fiber orientation distributions (FODs). This multi-tissue information was used to perform combined intensity normalization and bias field correction^69^. Liberal and conservative brain masks were used respectively for these two steps to mitigate the detrimental influences of imperfect masks in the respective processes.

Whole-brain tractography was performed as follows. A tissue-type segmentation image, intended for use of the Anatomically-Constrained Tractography (ACT) framework^70^, was constructed using a combination of the FreeSurfer `aseg` image and the results of FSL FIRST^23^. From this, a mask of the interface between GM and WM was constructed for the purpose of streamline seeding. Probabilistic tractography was performed using 2nd-order integration over Fibre Orientation Distributions (iFOD2)^71^. A total of ten million streamline seeds were drawn throughout the GM-WM interface, and generated streamlines were rejected if they failed to satisfy length constraints or the ACT priors^70^. This constant number of streamline seeds is an important requirement for computational tractability that provides a more robust upper bound on execution time across sessions.

### Structural connectivity: matrices

Tractography streamlines and parcellation images were next used to generate SC matrices quantifying various connectivity measures for each regional pair. To generate structural connectomes that are representative of whole-brain connectivity, we used parcellations that integrated both cortical and sub-cortical atlases. As described previously, structural connectivity matrices were pre-calculated for only a subset of all possible connectome configurations; the seven combinations of cortical and subcortical parcellations chosen—for which pre-computed out-of-the-box connectivity matrices are available—are summarized in Table 2. Nevertheless, the provided scripts and supplementary data (e.g. streamline endpoint coordinates and per-streamline metrics) enable connectivity reconstruction for any other possible combination of cortical and subcortical parcellation schemes desired.

**Table 2.** The seven combinations of cortical and subcortical parcellations for which structural connectivity matrices were precomputed, and their respective numbers of regions.

| Cortical atlas | Cortical regions | Subcortical atlas | Subcortical regions | Total regions |
| --- | --- | --- | --- | --- |
| Desikan Killiany | 68 | MSA - scale I | 16 | 84 |
| Destrieux | 148 | MSA - scale I | 16 | 164 |
| Glasser | 360 | MSA - scale I | 16 | 376 |
| Glasser | 360 | MSA - scale IV | 54 | 414 |
| Schaeffer 200 (7 networks) | 200 | MSA - scale I | 16 | 216 |
| Schaeffer 500 (7 networks) | 500 | MSA - scale IV | 54 | 554 |
| Schaeffer 1000 (7 networks) | 1000 | MSA - scale IV | 54 | 1054 |

Numerous measures of structural connectivity strength are available. The most common measure is the streamline count, which quantifies the total number of streamlines connecting region pairs. Alternatively, post processing algorithms can be applied to ensure that streamline counts better reflect the underlying white matter architecture. We provided per-streamline “weights” calculated by the SIFT2 method to estimate the Fiber Bundle Capacity (FBC) between regions^72^. Additionally, the average length of all streamlines between region pairs were computed to measure connection length. Finally, microstructural parameters (i.e. FA, MD, MO, S0, ICVF, ISOVF, and ODI) were averaged for each streamline trajectory to quantify the microstructural properties of connections.

Regarding the subset of connectome configurations for which we provide pre-computed structural connectivity matrices, for each of the combined parcellation schemes as shown in Table 2, matrices were constructed utilizing the following four metrics:

- Streamline count
- Fiber Bundle Capacity (from SIFT2)
- Mean streamline length
- Mean Fractional Anisotropy (FA)

For *all* structural connectivity metrics (ie. not only those for which matrices were pre-computed), the per-streamline quantitative metrics are provided alongside the locations of the endpoints of streamlines. This facilitates the construction of structural connectomes using any preferred combination of cortical parcellation, subcortical parcellation, and connectivity metric, utilizing the provided connectome generation scripts; this requires both minimal storage (only a pair of 3-vectors and a single floating-point value per streamline) and minimal additional computation (as it is the propagation of streamlines that incurs the greatest expense; assigning streamline endpoints to a parcellation is comparably simple). Considering all different parcellations and connectivity metrics, our code and data resources allow for user-friendly and time-efficient reconstruction of ∼1000 alternate structural connectivity matrices for a single subject. We further note that the data provided in this form are entirely compatible with the adoption of recent developments in the domain such as high-resolution connectomes that consider each surface vertex as its own parcel^73^, and the utilization of spatial smoothing of parcels to enhance reliability^74^, for which the relevant software tools are also provided.

### Computing resources

The whole pipeline was tailored for parallel execution on high-performance computing (HPC) clusters. Parallelization was implemented at the level of individual imaging sessions, with a separate computation job submitted for every session (∼ 45,000 parallel job submissions). While many of the underlying software tools are capable of executing multiple threads for a single processing job, this was not the case for *all* such tools, and therefore allocating a single CPU core per session was determined to yield the best CPU resource utilization. The maximal memory and wall time per job were empirically minimized to facilitate maximal parallelization on the HPC resource without compromising completion of jobs; this was chosen to be 4GB RAM and 6 hours execution time. The Spartan HPC resource provided by the University of Melbourne^75,76^ was utilized for this task, which was typically capable of executing 100–200 such jobs in parallel depending on external utilization.

#### Time requirements

In addition to the three primary computation steps of the pipeline, each computing job also involved downloading all required UKB data and uploading the resulting derivatives. While data download would typically only require minutes, UKB servers permit only 10 parallel downloads per user, and some jobs could hence experience considerable delays in accessing their requisite data; the wall time allocated for each job was therefore set in order to tolerate such delays without resulting in job failure.

Following data download, the approximate processing time required for each stage of the pipeline was as follows. The atlas mapping pipeline required 10 minutes. Mapping resting functional data required 5 minutes. The most time-consuming step was the structural connectivity reconstruction pipeline, which required 150 minutes. Finally, subsequent conversions and upload steps required 5 minutes. Overall, the complete pipeline required 3–4 hours to finish for a single imaging session.

In total, mapping connectomes across all UKB imaging sessions required ∼ 200,000 CPU hours (∼ 20 years) of computation to complete. This substantial time requirement could only be satisfied with the extensive use of HPC resources in parallel execution which reduced the overall required time to map these connectomes from years to months^75^.

#### Storage requirements

The storage requirements for the derivatives of the analysis pipelines are as follows:

- *Atlas parcellation pipeline*: Cortical and subcortical parcellations warped to native subject space, in addition to the non-linear warp from MNI to subject space (to facilitate reserarcher utilisation of other parcellations represented in MNI152 space), all stored in NIfTI format in a compressed archive, requires ∼100MB per session.
- *Functional connectivity pipeline*: The 28 time series (each unique cortical and subcortical parcellation as well as the global signal, as detailed in previous sections) are provided in comma-separated values (CSV) format. These files are provided as a compressed archive with a size of ∼50MB per session.
- *Structural connectivity pipeline*: The combination of parcellations, pre-calculated structural connectome matrices, streamline endpoint coordinates, and per-streamline quantitative metrics for construction of other connectome configurations, are provided as a compressed archive with a size of ∼75MB per session.

As a result, a total of ∼225MB of storage is required to store all derived data per session. The complete set of computed connectivity maps across all sessions occupies ∼10TB of storage.

## Data Records

The generated files are organized into three separate compressed archives for every UKB imaging session. Figure 6 provides a summarized list of files provided for a single session. Supplementary Figure S1 provides a more detailed list of all files. All data are made available via UKB data returns policy and will be accessible as new bulk files.

**Figure 6.**
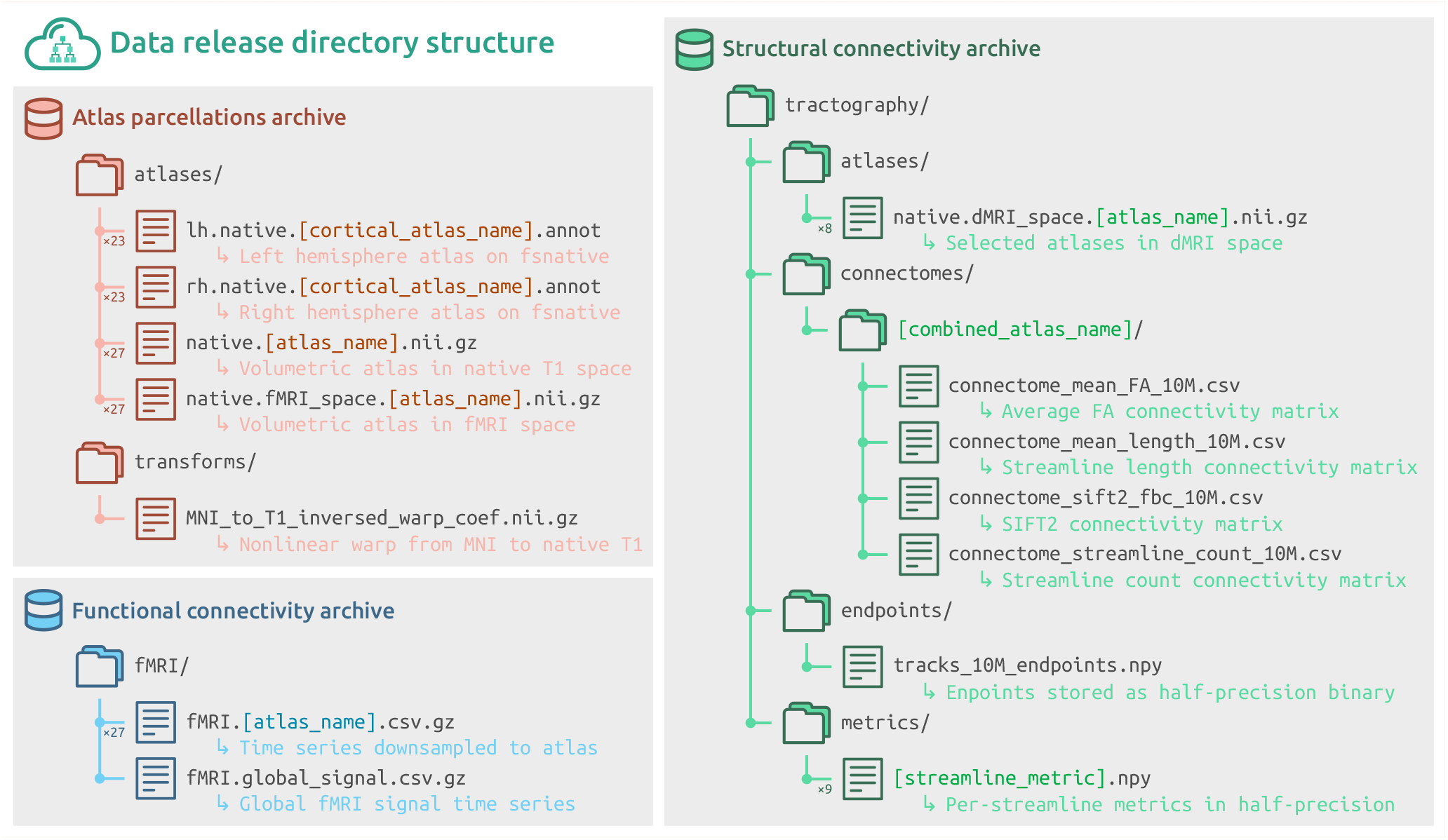
Summary of file archives comprising the connectome resource. Three separate compressed archives are provided for atlas parcellations (red), functional time series (blue), and structural connectivity data (green). Alternative configurations of provided data are summarized by providing a single informative placeholder (colored text in square brackets). A detailed list of all files is provided in Supplementary Information.

### Atlas data

The compressed bulk file of atlas data contains derivative parcellations in native volumetric space (`atlases/native. [atlas_name].nii.gz`) for all cortical and subcortical atlases used. Cortical atlases are also provided as parcellated surface data in the FreeSurfer “`fsnative`” space (`atlases/(rh|lh).native.[atlas_name].annot`). The warp image that can be used for nonlinear transformation from MNI space to native coordinates is additionally provided (`transforms/MNI_to_T1_inversed_warp_coef.nii.gz`).

### Functional data

The compressed bulk file of functional data contains time series are aggregated within the parcels of various atlases. The time series information is stored as a compressed file with comma-separated values, i.e. the `.csv.gz` format. For each atlas, all sampled time series are provided in a single bulk file (`fMRI/fMRI.[atlas_name].csv.gz`). In addition, the time series for the global signal is included (`fMRI/fMRI.global_signal.csv.gz`).

### Diffusion tractography data

The compressed bulk file of diffusion data contains:

- 28 pre-calculated connectivity matrices (all combinations of the combined atlases shown in Table 2 and the four primary connectivity metrics listed in the “Structural connectivity: matrices” section), with paths:(`tractography/connectomes/[cortical_atlas_name]+[subcortical_atlas_name]/connectome_[metric_name] _10M.csv`).
- Spatial locations of all streamline endpoints: (`tractography/endpoints/tracks_10M_endpoints.npy`). This facilitates assignment of pre-generated streamlines to any parcellation of interest. This file is stored in a half-precision floating-point format (16 bits) to reduce storage requirements with minimal loss of precision.
- Nine per-streamline quantitative metrics, with paths: `tractography/metrics/[metric_name].npy`. Metrics include streamline length, SIFT2 weights, and mean values of voxel-wise quantitative metrics along the streamline trajectories. These are also stored in half-precision floating-point format, with multiplicative factors of 1*e*^3^ and 1*e −* 3 applied to the mean MD and mean S0 metrics, respectively, to scale the magnitudes of floating-point values toward unity and therefore mitigate loss of precision.
- Statistics regarding tractogram generation (eg. why streamlines were terminated and why they were accepted or rejected).
- Statistics regarding the operation of the SIFT2 algorithm.

## Data Validation

All brain imaging data sourced as inputs passed the automated quality control (QC) evaluations by UKB^8^. We additionally computed various assurance metrics to assess the quality of the provided connectomic resource. These evaluations can be divided into i) QC metrics to probe data quality and ii) analyses of topological properties of brain networks, showing that our connectomes reproduce established findings of the network neuroscience literature.

### Quality control

We have computed an extensive set of connectomic QC measures. These metrics complement existing QC efforts provided by the UKB, and can be used to exclude low-quality or inaccurate connectomes. Figure 7 provides a summary of the provided QC measures.

**Figure 7.**
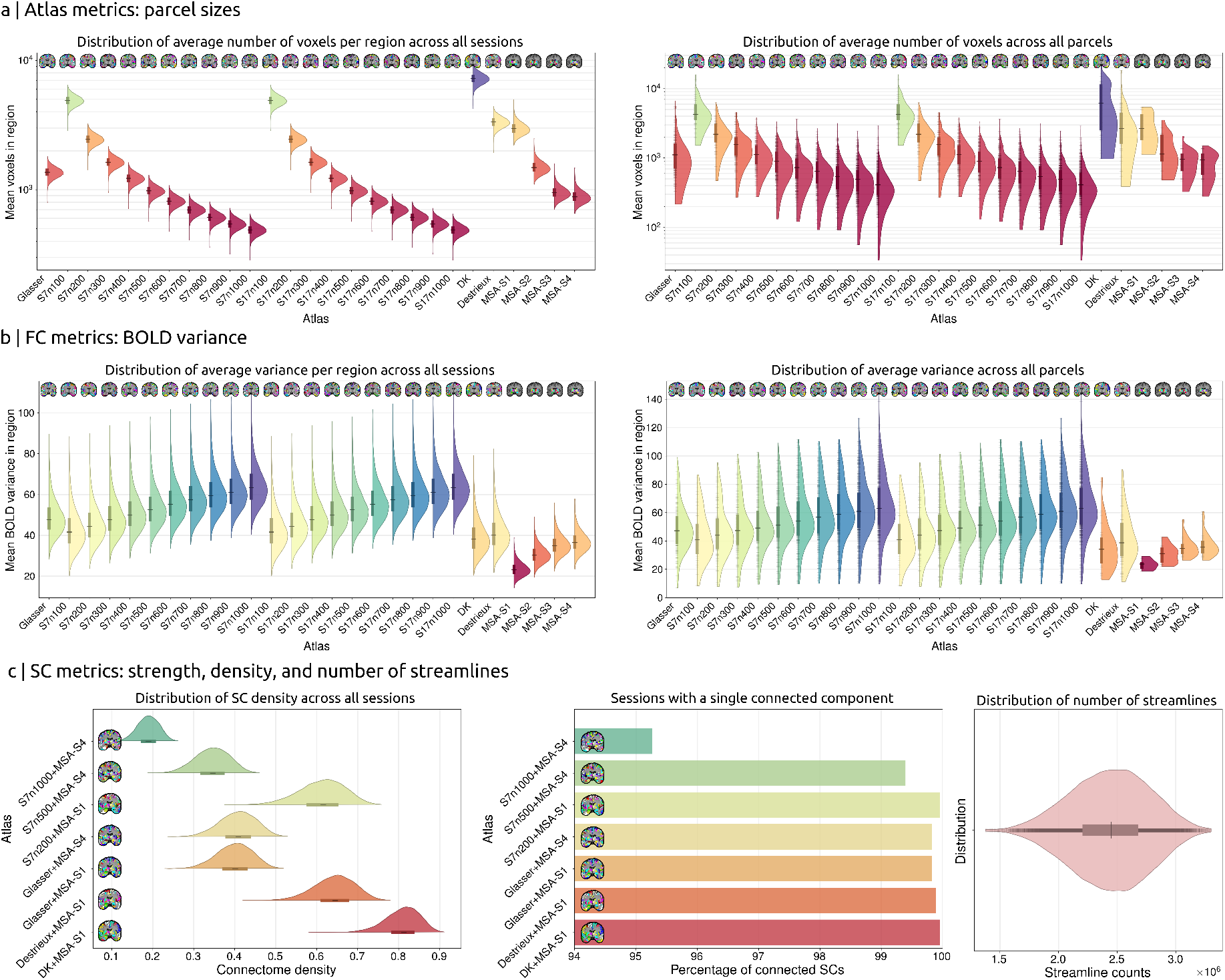
Quality control measures. (a) For atlas parcellations, the parcel sizes were quantified by the number of voxels within each region. Distributions of the number of voxels averaged across atlas regions (left) and across imaging sessions (right) summarize the trends in parcel size for all 27 cortical and subcortical atlases. (b) Similarly, distribution of BOLD signal variance is shown for all 27 atlases. (c) Summary of structural connectivity QC measures for connectome density, components, and reconstructed streamlines for all sessions.

Our QC measures serve as basic features that can be utilized to generate flexible data exclusion criteria tailored to specific research questions. For instance, cases where such metrics lie at the tail of the corresponding distribution could be excluded from analysis, or QC features could be matched between groups. We suggest that these QC features should be tailored to particular study aims, rather than being adopted blindly. For instance, one study may use the metric of total number of streamlines in the connectome to exclude individuals where tractography reconstruction is poor, or account for this effect as a confounding covariate; in contrast, another study aiming to assess the impact of a particular pathology on the structural integrity of the connectome may consider the total number of streamlines to be a variable of interest that should *not* be matched/regressed.

#### Atlas QC measures

For all volumetric parcellations (cortical and subcortical), the volume of every atlas region in an individual was computed as the total number of voxels assigned to each region. This measure can be used to investigate properties of the native atlas parcellations. Future studies may decide to exclude sessions or parcellations for which certain regions are not appropriately represented, e.g. if a region has no (or very few) voxels. In addition, these measures can optionally be used for normalization of particular structural connectivity matrices: since larger brain regions are more likely to be intersected by streamlines, one may choose to rescale connectivity based on regional volume^2,77^.

Figure 7.a shows a summary of this QC measure in the UKB sample. Violin plots on the left depict the distribution of voxel sizes averaged across all regions and plotted across sessions. Conversely, the plot on the right is averaged across sessions and depicts the distribution across regions. As anticipated, parcel volumes tend to become smaller as the granularity of the parcellation scheme increases. Furthermore, it indicates that a considerable degree of size variation exists between different regions of the same atlas; for instance, the largest regions of Schaeffer’s atlas with 1000 parcels are similar in size to regions from the Destrieux atlas with only 148 parcels.

#### Functional connectivity QC measures

For functional time series, the variance of the aggregated signal within each region is reported across all sessions and atlas regions. This regional measure of variance can be compared to the variance of the global signal to provide an estimate of signal quality^78^. However, it is important to interpret this information with caution, as the standard deviation of an fMRI signal cannot be equated to noise strength and is known to vary with aging and cognition^79,80^. Signal variance could still be used as a quantitative quality metric to filter out low quality scans.

Figure 7.b provides summary distribution plots of signal variance averaged across sessions and parcels. These summary plots show that relatively higher granularities (e.g. 1000 cortical regions compared to 100 cortical regions) tend to contain signals with larger variation. This is because large parcels sample fMRI over more voxels, which eliminates variance sources from localized effects and unstructured noise. These variance measures for QC could potentially be used as exclusion criteria for sessions in which a region has zero (or very low) variance. A retrospective evaluation of this QC measure indicated a limitation in UKB fMRI preprocessing pipelines impacting signal quality at the orbitofrontal cortex, which is known to be susceptible to BOLD signal loss^81,82^ (see Supplementary Information for further detail).

#### Structural connectivity QC measures

For structural connectivity, the number of streamlines reconstructed by tractography is reported as a measure of whole-brain tractogram reconstruction efficacy. Given that the number of seeded streamlines is constant for all individuals (10 million), a lower total streamline count indicates higher exclusion rates from streamline acceptance criteria (anatomical validity as determined by ACT and adequate length), which may indicate pathology, poor structural integrity, or low quality data. Another use of this feature is normalizing the connectivity matrices to construct connectomes with equal total strength, which can be more suitable for studying connection probability^83^. In addition, the number of connected components and total connectome density were computed for every structural connectivity matrix. This provides an additional QC feature, as connectomes are expected to form a single connected component and therefore disconnected connectomes could indicate either gross structural abnormalities or poorly reconstructed structural connectivity. Hence, future studies could exclude SC matrices with more than one connected component. As shown in Figure 7.c, SC forms a single connected component for most of the sessions (*>* 95% for the highest granularity, and *>* 99% for other granularities). Finally, the weighted nodal strength and binary nodal degree are provided, with a binarization threshold of one streamline. Figure 7.c summarizes the QC metrics extracted for structural connectomes.

### Properties of structural and functional connectomes

In this last section, we provide an initial exploration of the properties of our structural and functional brain networks. The following analyses sought to replicate previously established results in the network neuroscience literature and, as such, further demonstrate the quality and utility of our connectivity resource.

To this end, group-level structural and functional networks were constructed from connectivity data of 1000 randomly sampled individuals. We considered two parcellation combinations comprising a total of 68 (cortex: DK, subcortex: MSA-S1) and 554 (cortex: Schaefer 500, subcortex: MSA-S4) gray matter regions. FC was mapped using GSR and the pairwise Pearson correlation of regional BOLD signals. Group-level FC was computed as the average of functional networks across the 1000 subjects and group-consensus SC was inferred using consistency-based thresholding^84^.

Figure 8.a shows the resulting group average SC (blue) and FC (red) matrices for the two selected parcellation schemes. These visualizations align with previous literature^2,85,86^ and exhibit features that are consistent with typical connectomes, including i) distinct inter-hemispheric and cortico-subcortical boundaries, ii) a modular structure evident within each hemisphere, and iii) evidence of homotopic connections between the two hemispheres.

**Figure 8.**
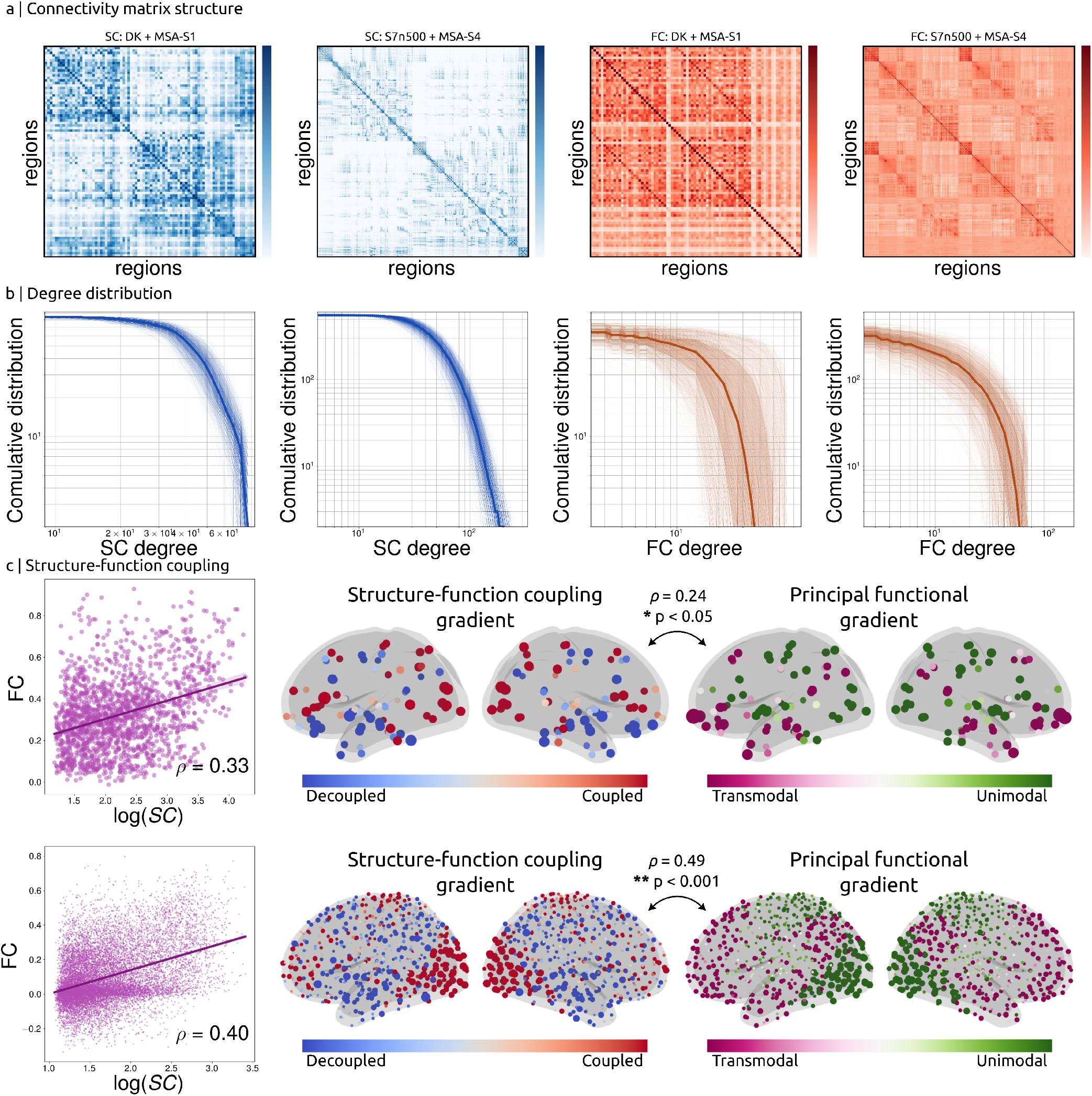
Properties of structural and functional brain networks. (a) Group-level connectomes were derived from 1000 random sessions. The matrices closely resemble standard brain connectivity characteristics with visual distinctions between the cortical hemispheres and the subcortex, local clusters of connectivity within each hemisphere, and strong homotopic connections forming diagonal strides. Results are presented for both SC (blue) and FC (red) matrices in two exemplar parcellations, with “DK + MSA-S1” containing the fewest and “S7n500 + MSA-S4” containing one of the greatest numbers of nodes. (b) Degree distributions of individual connectomes used to construct the group-level matrices. Thin lines indicate trajectories for individual session data; thick line presents the median; shaded regions indicate [25, 75] and [5, 95] centiles. (c) Structure-function coupling was quantified for both exemplar parcellations. Brain-wide coupling was assessed using Pearson’s correlation between FC and log(SC) across all edges (scatter plots). Node-wise coupling gradients were computed to project regions to a spectrum of decoupled (blue) to coupled (red) areas. These gradients were compared with FC-derived gradients of functional hierarchy in which regions are situated on a gradient from unimodal (green) to transmodal (purple) areas.

Next, we assessed the scale-free network property of the connectomes. We computed the degree distribution of the FC and SC matrices for all sessions by assuming respective binarization thresholds of *ρ* = 0.4 and 1 streamline, respectively. The degree distributions of the resulting matrices are presented in Figure 8.b. The corresponding degree distribution plots replicate previous findings of characteristics observed in structural and functional connectomes with a degree distribution that follows an exponentially truncated power law^86–91^.

Finally, we sought to reproduce previous findings of SC-FC coupling (Figure 8.c). The structure and function of the human brain networks are interrelated^92,93^. This relationship can be investigated by comparing the strengths of structural and functional network edges. We present data for three experiments in this regard:

1. *Structure-function correlation*: We assessed the degree of collinearity between FC strengths and the logarithm of SC strength (as quantified by streamline count). The expected positive correlation^94,95^ between the strength of structural and functional connectivity was observed for atlases with high (*ρ* = 0.4) and low (*ρ* = 0.33) parcellation granularities.
2. *Structure-function coupling gradient*: The degree of structure-function coupling is reported to vary across the brain, with certain regions exhibiting stronger coupling and others displaying relatively decoupled activity^96–99^. To evaluate structure-function coupling at the level of individual regions, we used a multilinear prediction approach^96^; in short, for each node, we estimated the FC strength to all other nodes via multilinear regression based on four measures of inter-regional distance and structural communication: (i) Euclidean distance, (ii) structural connectivity, (iii) shortest path length, and (iv) communicability^100^. The accuracy of model predictions (quantified by Pearson’s correlation) is indicative of local SC-FC coupling strength.
3. *Principal functional gradient*: In prior work, these local patterns of regional coupling have been reported to follow the functional organization hierarchy of unimodal to transmodal brain regions^97,98^. We thus computed the principal functional gradients by performing diffusion map embedding on group-level FC^101^ and evaluated its association with the SC-FC coupling gradient. Our results (Figure 8.c) successfully replicated the expected relationship between the coupling patterns and functional organization hierarchy at low (*ρ* = 0.24) and high (*ρ* = 0.49) parcellation granularities.

The evaluations of network properties presented in this section demonstrate that the connectivity matrices provided here reproduce well-established findings in brain connectivity research. This illustrates the high quality of this connectome resource and its potential to facilitate future connectomic studies in an aging population.

## Usage Notes

All data will be made available via UKB data returns policies to be accessible based on UKB material transfer agreements. Researchers can apply to access these data by filling out a UKB access application. Additional code and data (such as the label ordering of atlases) are made openly available in a publicly accessible git repository. All data are provided in formats that are readable in various programming languages. This enables use of several existing software packages for brain connectivity analysis such as the Brain Connectivity Toolbox (Matlab)^102^, bctpy (Python), and Nilearn (Python), as well as general network analysis tools such as NetworkX^103^.

## Code availability

All scripts used to perform computations described in this manuscript (eg. generating atlas parcellations, aggregating BOLD time series, tractography, computing structural connectivity, etc.) are made publicly available in the git repository served at github.com/sina-mansour/UKB-connectomics. Additional scripts used to perform QC evaluations are also provided in this repository. Furthermore, sample scripts to read the generated data into various programming languages are made available to facilitate uptake of this resource by the community.

## Acknowledgements

This project was funded by CS’s Early Career Researcher Grant from the University of Melbourne. The data analysis was supported by Spartan High-Performance Computing infrastructure^75,76^, and dedicated data storage solutions provided by the Research Computing Services at the University of Melbourne. RS is a fellow of the National Imaging Facility, a National Collaborative Research Infrastructure Strategy (NCRIS) capability, at the Florey Institute of Neuroscience and Mental Health.

## Author contributions statement

**S.M.L**.: Pipeline development, Software, Data curation, Formal analysis, Writing - original draft, Writing - review & editing. **M.D.B**: Pipeline development, Writing - original draft, Writing - review & editing. **R.S**.: Pipeline development, Software, Writing - original draft, Writing - review & editing. **A.Z**.: Supervision, Pipeline development, Software, Writing - original draft, Writing - review & editing. **C.S**.: Supervision, Funding acquisition, Pipeline development, Writing - original draft, Writing - review & editing.

## Competing interests

The authors declare no competing interests.

## Supplementary Information

### Comprehensive directory tree

A summarized directory tree was provided in the main text to list all released data. Figure S1 provides an extensive list of all files provided in the compressed archives of a session.

**Figure S1.**
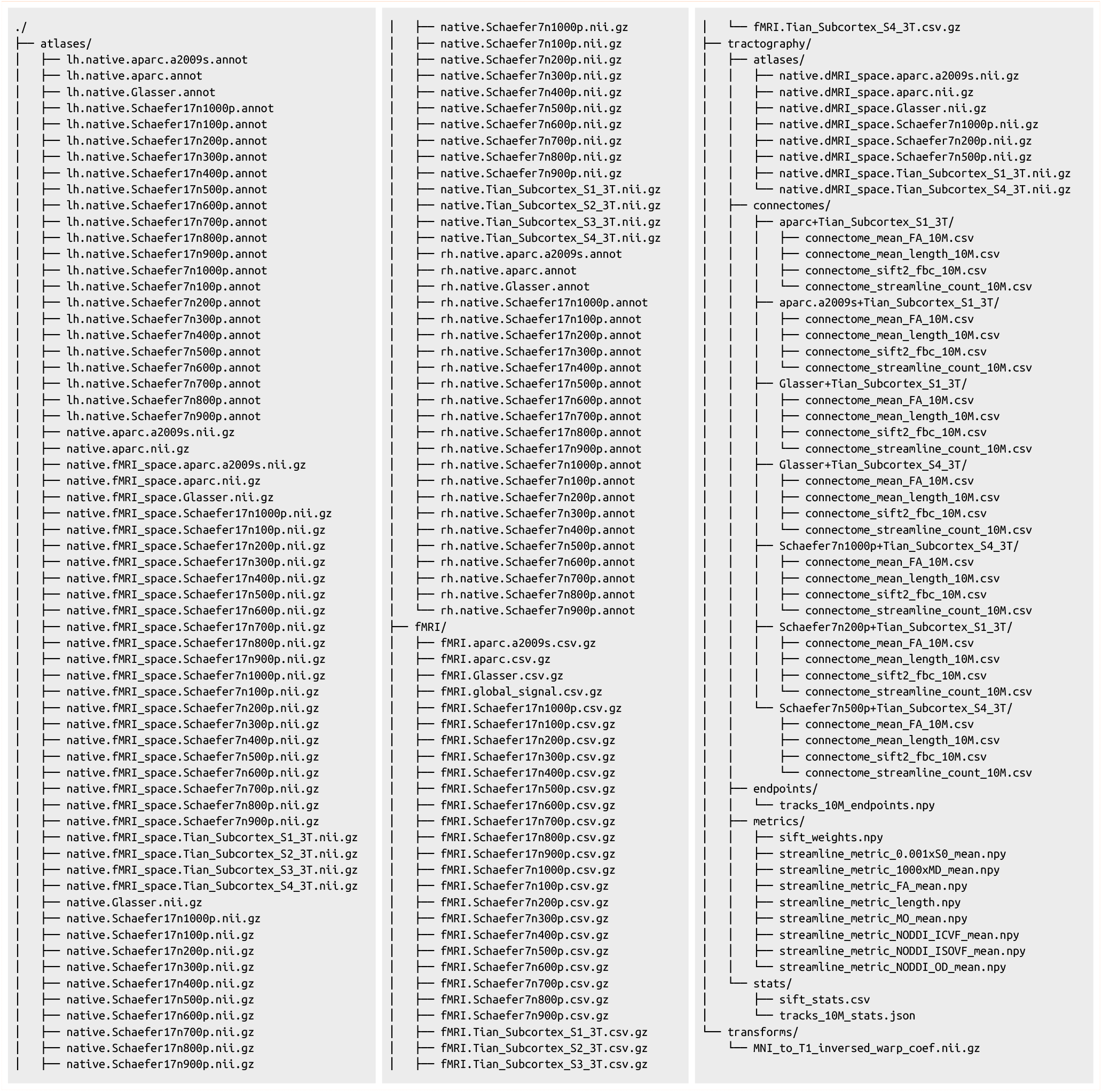
Complete list of all files included in provided archives. Three separate compressed archives are provided, for each of atlas parcellation (↪`atlases/*`), functional time series (↪`fMRI/*`), and structural connectivity (↪`tractography/*`) pipelines.

### Resting-state signal loss

The BOLD signal was collected using a gradient echo EPI acquisition which can experience geometric distortion as well as signal loss in regions of strong susceptibility^81^. Computing a brain mask from empirical fMRI data, using any method based on image intensity, may lead to exclusion of such regions. All voxels outside of this brain mask were explicitly zero-filled in UKB fMRI preprocessing pipeline^8^. For a region that can be clearly delineated on an effectively undistorted anatomical image, but its entire volume resides outside of the fMRI brain mask, the aggregate fMRI signal within that region would consequently be zero-filled. These cases can be identified by computing the QC measure of regional signal variance. Hence, signal variance can be used as a proxy to assess signal loss due to exclusion of susceptible regions.

To evaluate the cases in which UKB preprocessing led to such problems, we counted the number of sessions for which a variance of zero (constant signal) was reported for a brain region. This information was then sorted to pinpoint the regions with highest detrimental impact of signal loss. Table S1 provides a list of percentages of signal loss for 100 mostly affected brain regions (across different atlases) sorted based on signal loss severity. This indicated a pattern of severe signal loss within the orbitofrontal cortex (OFC). The signal loss severity was higher for higher granularity levels such that the cortical atlases with 1000 regions had signal loss in more than a third of the cohort. However, the severity was considerably lower for lower granularities such that atlases with fewer than 700 regions had signal loss in less than 5% of the cohort.

We hereby report this issue as a limitation of the fMRI acquisition and preprocessing and suggest either i) conducting the analysis at lower granularities (especially if OFC is of interest), or ii) excluding the impacted regions of OFC in cases where the analyses are to be conducted on higher granularities. Finally, to verify that signal loss was due to impacts of preprocessing on OFC and not a result of the connectivity mapping pipeline manual visual inspections were conducted for a handful of sessions. Figure S2 provides an example that shows a complete lack of signal in the OFC parcel (`Schaefer7n1000p:7Networks_LH_Limbic_OFC_6`) due to zero-filling leading to a zero variance QC metric.

**Figure S2.**
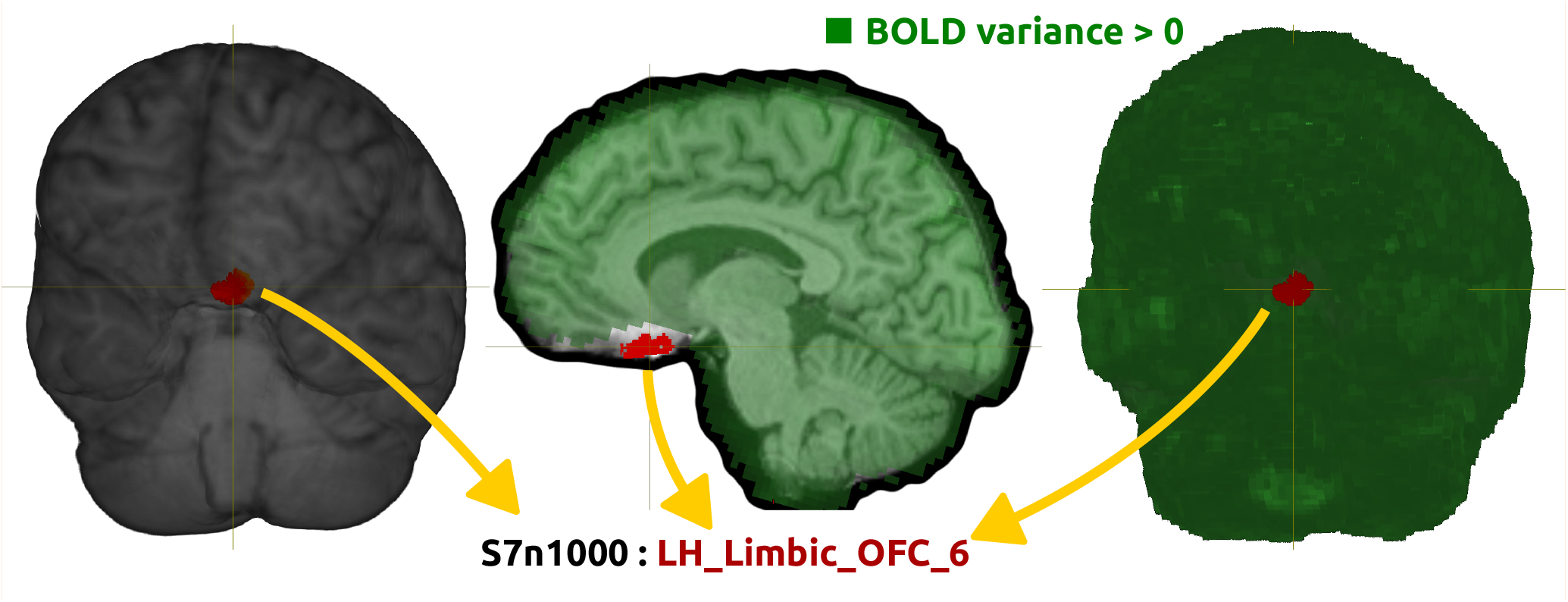
Illustration of signal loss near certain OFC regions for a single session. The `LH_Limbic_OFC_6` regions from `S7n1000` atlas is highlighted (red) on a 3D rendered T1 brain [left]. A sagittal slice of the brain showing the same parcel (red) along with a highlighted map of regions within the rfMRI brain mask (green) for which signal variance is positive [middle]. A similar 3D render showing the particular OFC parcel falling outside of the fMRI mask [right]. The images indicate that the preprocessed fMRI signal fails to cover certain cortical gray matter regions belonging to the OFC.

**Table S1.**
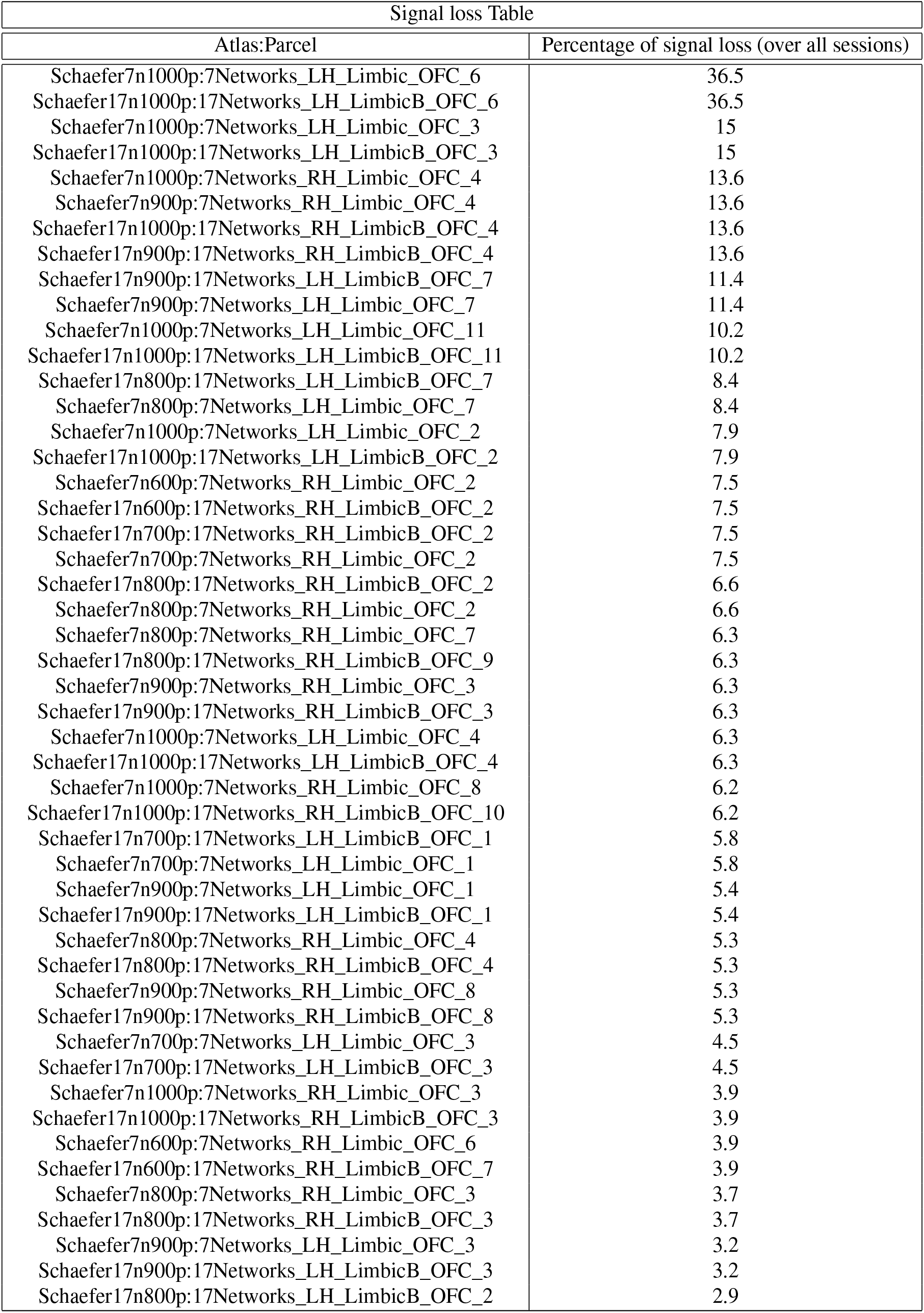
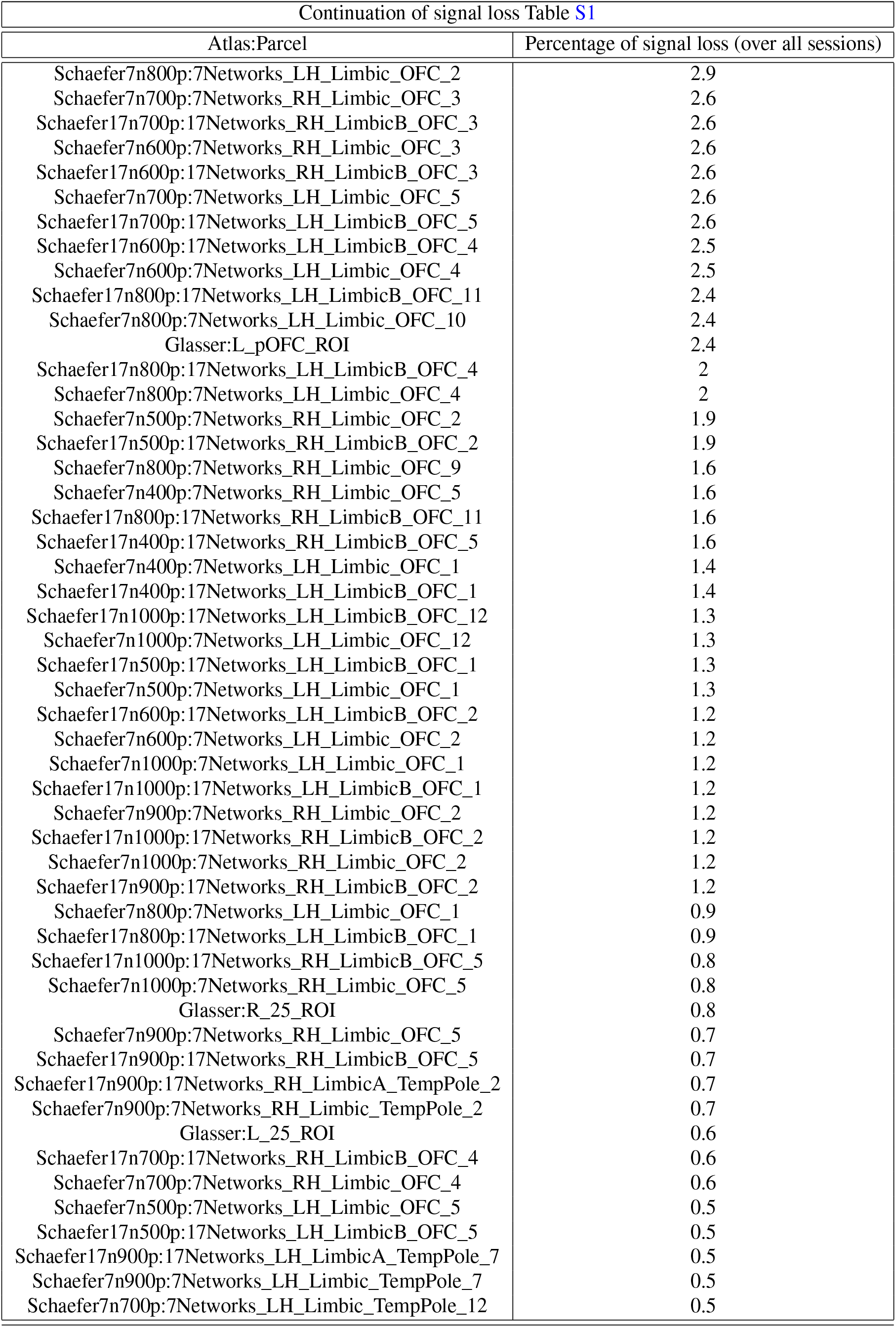
One hundred regions that were most severely affected by BOLD signal loss. The percentage of sessions for which zero variance was recorded in a certain region was quantified. This quantifies the frequency by which a region falls outside of the fMRI brain mask due to effects of signal loss.

## Notes

https://github.com/sina-mansour/UKB-connectomics

### Competing Interest Statement

The authors have declared no competing interest.

